# Human gut microbiota subspecies carry implicit information for in-depth microbiome research

**DOI:** 10.1101/2025.02.05.636567

**Authors:** Matija Trickovic, Silas Kieser, Evgeny M. Zdobnov, Mirko Trajkovski

## Abstract

Microbial strains from same species can have distinct functional characteristics owing to their different gene content. As the highest resolution, strains are mainly host-specific, thus obscuring unbiased associations, and hindering deductive research. Here, we comprehensively define the human gut microbiota at consistently-annotated subspecies resolution in an unbiased, cohort-independent manner, and demonstrate that we can generalize across distinct populations worldwide while maintaining specificity and improving interstudy reproducibility. We developed panhashome, a sketching-based method for rapid subspecies quantification and identification of genes that drive the intraspecies variations, and showed that subspecies carry implicit information undetectable at species level. By meta-analysis of colorectal cancer (CRC) datasets, we identified disease-associated subspecies whose sibling subspecies or species are not. Subspecies-based machine-learning CRC diagnostic algorithm outperformed species-level methods by leveraging the unique subspecies-level information. This subspecies catalogue allows identification of genes that drive the functional differences between subspecies as fundamental step in mechanistically understanding microbiome-phenotype interactions.

## Introduction

The development of advanced computational tools enables rapid and precise quantification of microbial abundances mainly by using custom references^1,2^, with the majority of them being annotated to species level. Strains within the same species can notably differ in their gene content, and these intraspecies variations can drive distinct functional characteristics between strains belonging to the same species^3^. Conflating different strains can obscure associations with a condition or a treatment, thus hindering deductive research. One of the main challenges in establishing reproducible causal links between the microbiome and health is the limitation of current analysis methods to species level due to these intraspecies variations. The bacterial strains, as the highest resolution, are mostly host-specific^4^. Strain definitions often differ between fields^3^. In metagenomics, a strain is usually defined as a single genome or a metagenome-assembled genome (MAG), which translates to a very high number of total strains that would further grow with every additional detected single nucleotide variations (SNVs), and which can be decreased with thresholding the distance between genomes. Recent developments render it possible to analyze a fraction of the bacterial species within a microbiome at the strain-level resolution^5,6^. However, that is usually done for one or a couple of species, and most existing algorithms only focus on the dominant strain within a sample^4,7–9^. Although it is possible to differentiate strains based on SNVs, current tools have several drawbacks. Detangling species variability based on calling SNVs directly from sequencing data^4,10,11^ defines new subspecies groups for every analyzed cohort, thus disabling formulation of generally applicable conclusions, albeit acknowledging the intraspecies variability. Systematically and comprehensively quantifying the microbiota at the strain level is therefore unsuitable. Accordingly, linking strain identities to strain-specific functions is difficult, contributing to challenges in investigating the underlying mechanisms of microbiome-disease interactions. These issues in microbiome research suggest that both species, and strain-level microbiome profiling are suboptimal for differential abundance meta-testing across samples and studies, highlighting the need to enhance the resolution^12–17^. Recent work addresses the level between species and strains – subspecies resolution as narrowest taxonomic group within bacterial species that show functional, phenotype-specific differences towards its sibling subspecies (i.e. subspecies delineated from the same species), in one^18,19^, or a limited range of selected parental species^20^ (i.e. species within which the sibling subspecies are delineated). Such resolution would bypass the inheriting lack of reproducibility between subjects and studies at a strain level, but would optimally account the intraspecies heterogeneity. In light of the above, generating a consistently annotated catalog of the human gut microbiota on a subspecies level is of critical need in microbiome research, which would enable the identification of subspecies with underappreciated roles in human health and disease.

In this study, we developed the first comprehensive <u>hu</u>man gut <u>m</u>icrobiota <u>sub</u>species (HuMSub) catalogue by defining the human microbiota at subspecies resolution in an unbiased and cohort-independent manner. We demonstrate that at this resolution, we can generalize across different and distinct populations worldwide while still maintaining specificity, and show that subspecies carry novel implicit information not present at species level. We challenge this concept using meta-analysis of all available colorectal cancer patient datasets, where we identify subspecies associated with the disease whose sibling subspecies or parental species are not. Since sibling subspecies share a large portion of their genome, we establish a unique setup for fast and easy subspecies quantification, as well as identification of specific genes that drive the subspecies variations and link them to the host phenotype. We develop a subspecies-based machine-learning approach demonstrating that the novel layer of information provided at subspecies level consistently outperforms the machine-learning algorithms that rely on species resolution. Identifying specific disease-associated microbial genes and, hence, functions is fundamental in understanding the underlying mechanisms of microbiome-disease interactions. This catalogue, and the workflows we developed allow identification of the specific microbial genes that drive the sibling subspecies differences, thus providing basis for mechanistic, and potentially causative explanations for their differential associations with diseases. The HuMSub catalogue contains a comprehensive collection of consistently annotated subspecies of the human microbiome to date, setting the ground for analysis of new and reanalysis of the existing datasets at an unprecedented depth.

## Results

### Subspecies from the HuMSub catalogue are ubiquitous across the bacterial domain and are directly quantifiable

We used the genomes from *HumGut*^21^, the most comprehensive catalog of bacterial genomes from the human gut at the time of starting the project. HumGut was built using the UHGG catalog^22^, with additional assembled and reference genomes. While the genomes initially met the medium quality standard^23^, we applied more stringent quality filtering based on GUNC^24^ and BUSCO^25^ to further remove remaining contaminated and incomplete genomes (Figures S1A and S1B). For subspecies clustering, we first created sketches of coding sequences predicted in all genomes using Sourmash^26^. We did this with the expectation that variants in coding sequences would lead to identification of distinct subspecies phenotypes instead of hard-to-interpret intergenic variation. For species with more than three genomes, we then calculated the pairwise distances and clustered them in an unsupervised manner using the Prediction Strength metric^27^ with a cutoff of 0.8 to select the optimal number of clusters. We confirmed that the larger input species with more than 50 genomes are indeed homogeneous, as recommended for the Prediction Strength metric (Figures S1C, D).

After the quality filtering and clustering, we kept 225918 genomes represented across 3483 species. Using the above-defined approach, we generated the HuMSub catalogue (Figure 1A), where we identified a total of 5361 subspecies belonging to 977 species, revealing that 28% of species contain previously routinely neglected subspecies variations (Figures 1B and S1E). Notably, the clustering protocol didn’t induce a taxonomic bias, as ratios of phyla remained similar between species and subspecies levels (Figure S1F). 70% of subspecies contained more than one reference genome (Figure S1G), suggesting that the clustering approach effectively generalizes across the bacterial genomes.

**Figure 1:**
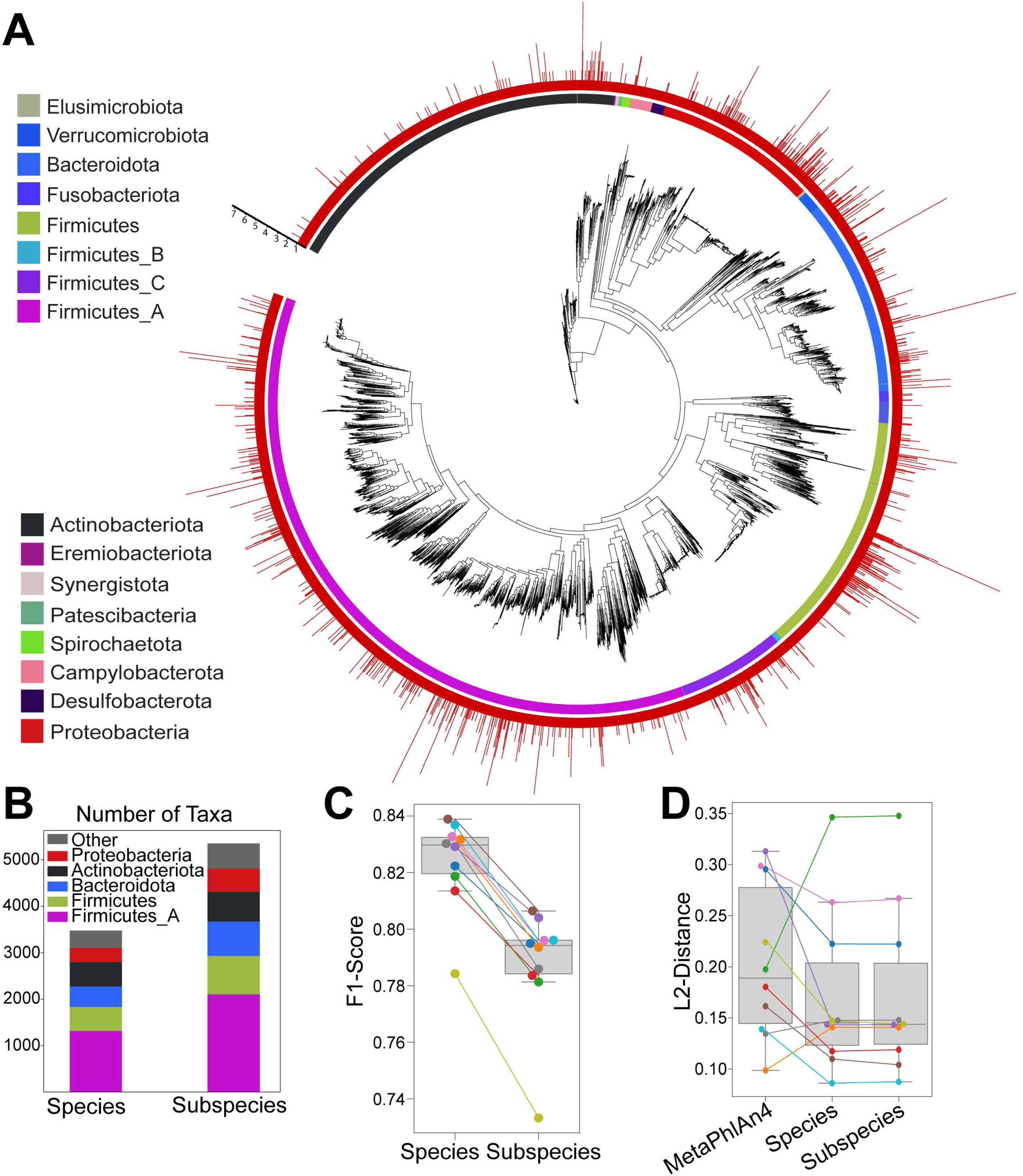
Subspecies from the HuMSub catalogue are ubiquitous across the bacterial domain and are directly quantifiable. (A) Phylogenetic tree of 3483 species. The inner color ring represents phyla attribution, and the height of the red bars shows the number of detected subspecies. (B) The number of different taxonomic categories denoting the five largest phyla. (C) Boxplot of F1-score distribution at species and subspecies level. (D) Boxplot of L2-distance distribution at species and subspecies level. MetaPhlAn4 performance is included only in L2-distance calculation since it defaults to lower taxonomic levels if a higher one can’t be determined, which would result in an artificially low F1-score in this benchmark setting. The colors in (A) correspond to (B) and represent phyla, while the colors in (C) correspond to (D) and represent same samples.

Quantifying subspecies was previously challenging due to the high genetic similarity and large genomic segments that are shared among strains within the same species. While methods utilizing SNVs can quantify strains and subspecies, they only offer an indirect calculation of their abundance based on the overall species abundance. To overcome these issues, we established a method for direct quantification of the previously defined subspecies from metagenomic sequencing, for which we developed a custom-defined concept of subspecies-specific “panhashome”, as a collection of hash values specific to each subspecies. Specifically, we first sketched all genomes in the species using Sourmash^26^. Then, we chose the hashes that appeared in at least 20% of the genomes of a particular subspecies, but in less than 5% of the genomes from different subspecies (Figure S2A). We constructed a subspecies quantification benchmark to validate the newly established quantification approach. We simulated ten metagenomic samples and assessed the Euclidian (L2) distance, which measures the difference between ground truth and quantified abundances, and the F1-score as a rate of false positives and false negatives, on both species and subspecies resolution. Additionally, we tracked the computational resources required for the quantification. To help interpret the performance metrics, which could be influenced by the complexity of simulated microbiome samples, we included MetaPhlAn4^1^ as a computational tool for species-level profiling commonly used in the field. Since there is no possibility of including custom references in MetaPhlAn4, we used it only at the species level. The median F1-score and L2 distance were 0.794 (Median absolute deviation (MAD): 0.009; Standard error of mean (SEM) 0.006) and 0.144 (MAD: 0.032; SEM 0.026), respectively, suggesting a very high performance of our approach (Figures 1C and 1D) for both species- and subspecies-level profiling. Both runtime and total memory use were markedly reduced when using our sourmash-based approach compared to MetaPhlAn4, showing that the quantification of subspecies is both faster and less memory intensive compared to previous state-of-the-art species-level quantification (Figures S2B and S2C). The high performance of the sourmash-based approach is in agreement with the LEMMI independent benchmark^28^.

To further validate our quantification approach on novel genomes, we simulated a new set of ten metagenomic samples using genomes uploaded to RefSeq after the publication of the HumGut catalogue. We kept the ones whose NCBI taxonomic IDs were originally contained in the catalogue and further quality-filtered them. To annotate them with subspecies information, we used the similarity of their sketches to the subspecies-specific ones and kept genomes with detected similarity > 0.8 (see Methods for details). This benchmark enabled us to assess the quantification performance in real-world use. We obtained similar results, with a median F1-score of 0.777 (MAD: 0.019; SEM 0.008) and L2 distance of 0.151 (MAD: 0.042; SEM 0.030), further indicating high performance on real-world data (Figures S2D and S2E).

### Global biogeographical subspecies distribution

The human gut microbiome varies across different populations, even among cohorts otherwise similar in phenotype^29–31^. The variability increases by enhancing the taxonomic resolution, to the extent that most strains are not only population, but even host-specific^4^. This hinders deductive research and prevents strain level from becoming widely used in metagenomic quantification.

Given that subspecies offer an increased resolution compared to species and might face similar limitations, we explored its potential utility for meaningful global-scale microbiome research. We searched 5272 publicly available human metagenomic samples with country information for the presence of subspecies using Mastiff^32^. By grouping the samples to a continent level, we found that 62% of the quantified subspecies are found in at least three of the four analyzed continents, while only 2.5% of included subspecies showed zero prevalence (Figure 2A). The African continent showed the highest number of specific subspecies (Figure 2B), which could be explained by the different host lifestyles. The three African countries: Madagascar^17^, Cameroon^33^, and Tanzania^34–36^, indeed show previously neglected Africa-enriched microbiome, further confirmed by a recent study^37^, with subspecies that were shared across those countries (Figure S3A).

**Figure 2:**
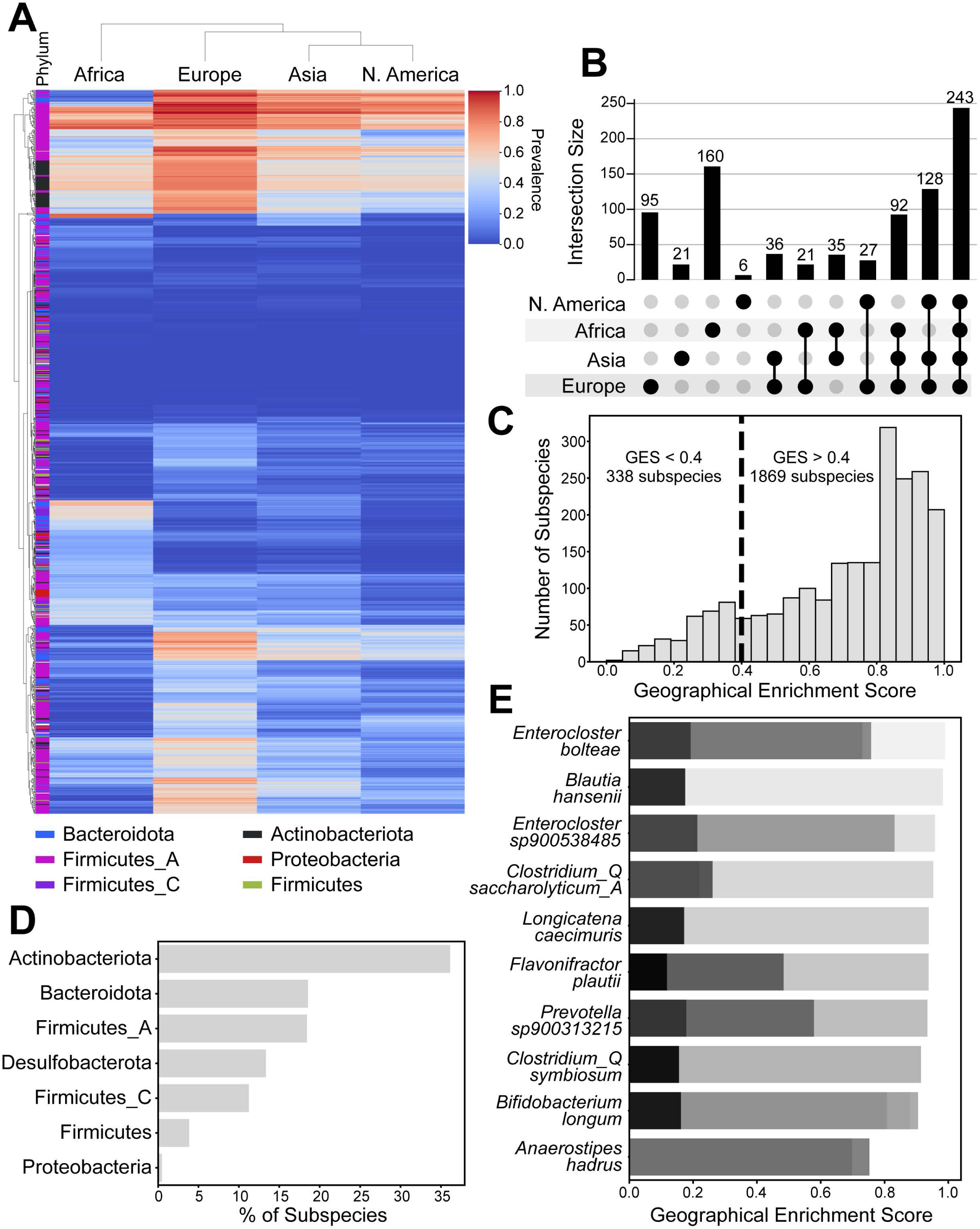
Global biogeographical subspecies distribution. (A) Clustered heatmap showing the prevalence of subspecies from the six biggest phyla. (B) UpSet plot showing the number of detected subspecies shared by different continents. (C) Histogram showing the distribution of the Geographical Enrichment Score (GES) for all analyzed subspecies. The vertical dashed line, at GES=0.4, denotes the threshold for geographically restricted subspecies. (D) Barplot showing the percentage of geographically restricted subspecies (GES < 0.4) for the top five analyzed phyla. (E) Stacked barplot of GES values of the top ten species where at least one subspecies is geographically restricted (GES < 0.4).

To better summarize the biogeographical distribution of the subspecies, we defined a single metric that helped us survey their enrichment. We compared the prevalence of subspecies between all countries and weighted the median difference with adjusted *P*-values to get a Geographical Enrichment Score (GES ϵ [0,1]) for each subspecies (see Methods for details). Calculated as such, GES represents the level of geographical restriction of a subspecies, where GES=1 shows no restriction, and GES=0 demonstrates high country-level specificity. To validate the metric, we confirmed its correlation with median adjusted *P*-values (Figure S3B) for each subspecies obtained through cross-country pairwise comparisons. From there, we concluded that GES=0.4 effectively distinguishes country-specific versus wide-spread subspecies. We then used this threshold to split all subspecies into groups of high and low specificity. By looking at the distribution of median differences in prevalence obtained through the same comparisons as *P*-values, we found that subspecies characterized with low specificity showed minimal median differences in prevalence compared to highly specific ones with a much broader distribution (Figure S3C). Based on this threshold, 1869 subspecies were shared for many of the analyzed countries, and 338 subspecies were found specific to a certain region (Figure 2C). These country-specific subspecies accounted for a high percentage of Actinobacteria, Bacteroidota, and Firmicutes phyla (Figure 2D). Firmicutes phylum contains the highest total number of restricted subspecies (Figure 3SD), in agreement with a previous study^20^. By looking at the species whose subspecies showed the highest difference in GES, we identified examples where one subspecies is geographically restricted while others aren’t (Figure 2E). This represents novel information that would be missed when analyzing at both species and strain resolution. The highest difference showed *Anaerostipes hadrus*, where one of its subspecies was highly prevalent in Great Britain, Denmark, Sweden, Korea, and the Netherlands and almost completely absent in populations from Canada, India, and Madagascar, while its sibling subspecies are lowly but similarly prevalent everywhere. Collectively, these data suggest that unlike strains, the subspecies within the HuMSub catalogue generalize across different and distinct populations worldwide while maintaining specificity.

### Subspecies contain implicit information not available at species resolution

Colorectal cancer (CRC) is the third most common cancer responsible for the second-most cancer-related deaths worldwide^38^. To assess the practical use of the HuMSub catalogue, we used microbiome samples from seven CRC studies^39–45^, totaling to 555 CRC samples and 530 healthy controls, and quantified them at the subspecies level. Out of the 2800 identified subspecies, we found in total 218 significantly associated ones (abs(random effect size) > 0.2; FDR < 0.1). Out of those, 104 were instances where at least one subspecies was significantly associated with CRC, while at least one sibling subspecies was not (Figures 3A, marked in blue, S4A and Table S1). As a proof of principle, we detected *Fusobacterium animalis*, previously associated with CRC development^46^, with two sibling subspecies, where only *001002* is significantly increased in cancer samples (Figure 3B). A recent study^47^ focusing entirely on this species experimentally demonstrated the existence of two clades, where only one populates the tumor niche and contributes to intestinal adenomas. Similarly, *Porphyromonas asaccharolytica*, also described as a contributor to CRC^48^ showed subspecies *001002* as one of the highest CRC-associated subspecies, unlike its sibling subspecies *001001*, which didn’t show association with CRC (Figure S4B).

**Figure 3:**
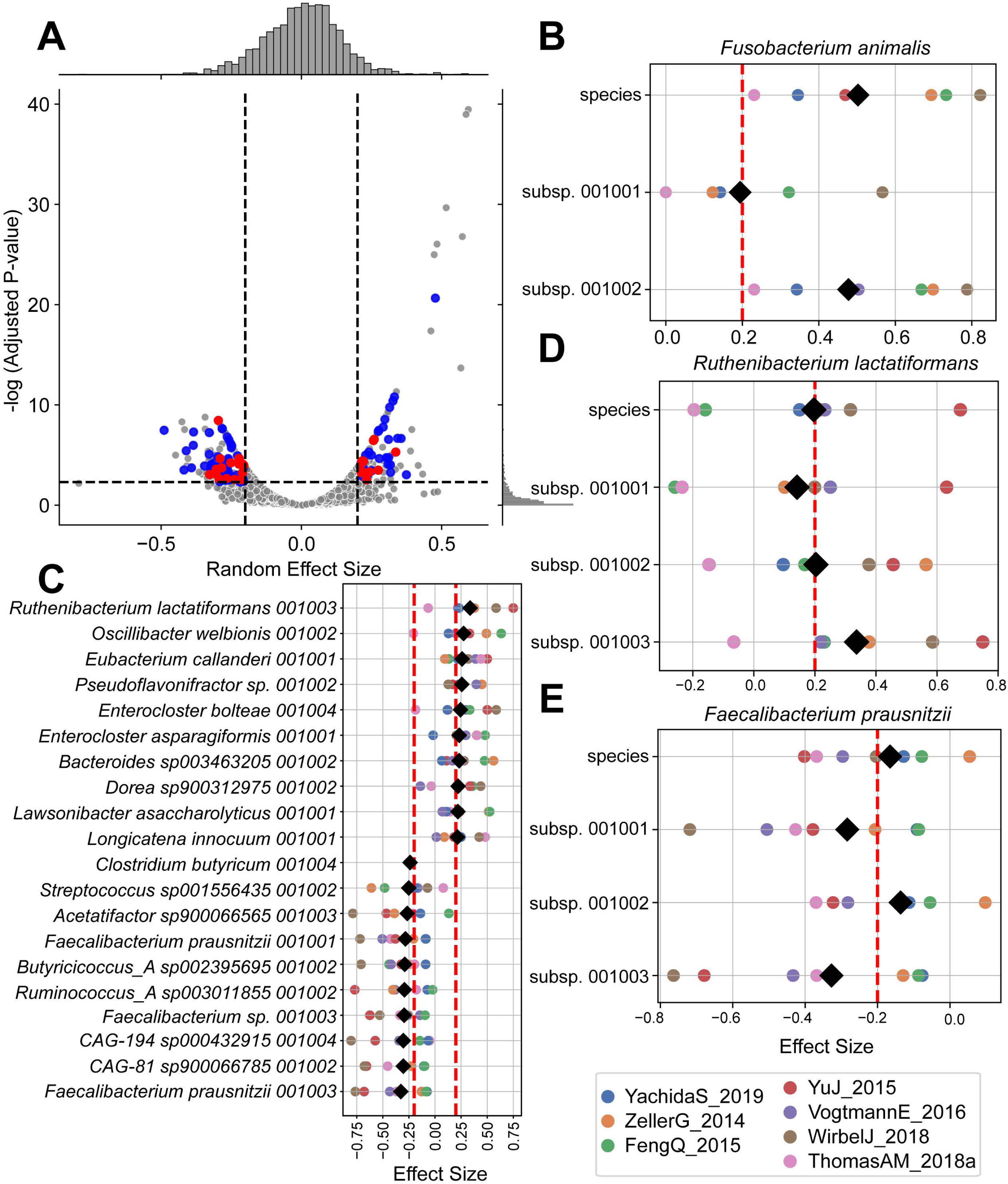
Subspecies contain implicit information not available at species level. (A) Vulcano plot showing the Random Effect Size and adjusted *P*-value at the subspecies level. Blue dots represent significantly associated subspecies whose sibling subspecies aren’t associated. Red dots represent significantly associated subspecies whose parental species aren’t associated. (B) Forest plot of effect sizes and random effect sizes for *Fusobacterium animalis* at species and subspecies levels. (C) Forest plot of effect sizes and the random effect sizes of the top ten subspecies positively and negatively associated with CRC, while their parental species isn’t (Red dots from panel A). (D and E). Forest plots of effect sizes and random effect sizes for *Ruthenibacterium lactatiformans* (D) and *Faecalibacterium prausnitzii* (E) species and their subspecies. In B-E color dots represent effect sizes estimated for each study individually using standardized mean difference, while black diamonds represent random effect sizes. Species and subspecies are significantly associated if abs(random effect size) > 0.2 and adjusted *P*-value < 0.1.

On the other hand, 28 subspecies were associated with CRC where the parental species didn’t show association with disease, representing 26.9% of previously discussed associations (Figures 3A, marked in red, 3C and Table S2). Among these, the subspecies *Ruthenibacterium lactatiformans 001003* was increased, while subspecies *001001* and *001002*, together with the species level, were not significantly different in the samples from CRC subjects compared to those from healthy controls (Figure 3D). Similarly, the subspecies denoted as *UBA1691 sp900544375 001002* was increased in CRC, while its sibling subspecies *001004* and the species level of this bacterium were not (Figure S4C). Contrary to this, some subspecies were less abundant in samples from CRC subjects. For example, *Faecalibacterium prausnitzii* is a species described as beneficial in ameliorating CRC^49,50^. In the meta-analysis, two of its subspecies were decreased in samples from subjects with CRC, while the species level itself wasn’t (Figure 3E). This indicates that not only does subspecies level provide new insights into statistical and causal associations with a disease, but that they could also improve reproducibility and to some extent explain discrepant results between studies. To further examine this possibility, we used univariate statistics for differential abundance testing within each study individually, both at the species and subspecies level, and found 31 species whose subspecies were associated with CRC in more studies compared to the species level alone (Figure S4D). This demonstrates that subspecies contain a layer of information fully neglected or missed when analyzing the microbiome on a species level and that the subspecies are better suited for cross-study comparisons, providing associations with improved replicability between cohorts.

### Subspecies are better predictors of colorectal cancer

To further investigate the usability and practical importance of the findings that subspecies carry more informative gut microbiome profile stratification than species, we used machine learning to train classifiers for distinguishing samples from CRC and healthy subjects. We used a Light Gradient Boosting Machine (LGBM)^51^, an ensemble-type model designed to work well with sparse and high-dimensional data. To directly compare the information content of species vs. subspecies inputs, we trained same-structured models on two different levels: MetaPhlAn-based species-level genome bins (SGBs) abundances and subspecies-level profiles quantified using the HuMSub catalogue. To assess their performance, we either pooled all samples together (cross-validation), or used the Leave-One-Dataset-Out (LODO) approach^52^. The former allows the model to train on some samples from all studies, thus enabling potential per-study adjustments. On the other hand, with the LODO approach, the model is trained on all studies except one, which is then used entirely for testing.

We used the samples from all seven up-to-date reported studies as in the meta-analysis. Importantly, species-level models achieved median performance similar to the current literature in this kind of setting (AUROC=0.79)^44^, establishing a credible baseline. During the cross-validation, subspecies-level models mildly outperformed species-level ones (Figure S5A). Strikingly, by employing the LODO approach that simulates real-world scenarios, we found that subspecies-level models (median AUROC=0.838, exceeding 0.890 for select datasets) consistently outperformed species-level ones (median AUROC=0.785) for all 6 studies (Figure 4A), except when testing on VogtmannE_2016^45^, which similarly resulted in lowest prediction rates as previously reported at species level^44^. These results suggest that subspecies resolution inherently contains novel information, which is more transferable between studies, explaining the increase in performance. Similarly, by training the models on only one of the studies and testing on all others, we observed that in two-thirds of study-to-study comparisons, subspecies-level classifiers outperformed the species-level ones (Figure S5B). Together with an increase in negative predictive value (NPV), an important metric for screening-type tools (Figure S5C), the increase in the performance of subspecies-level models suggests their superior predictive power, making them more suitable for clinical use.

**Figure 4:**
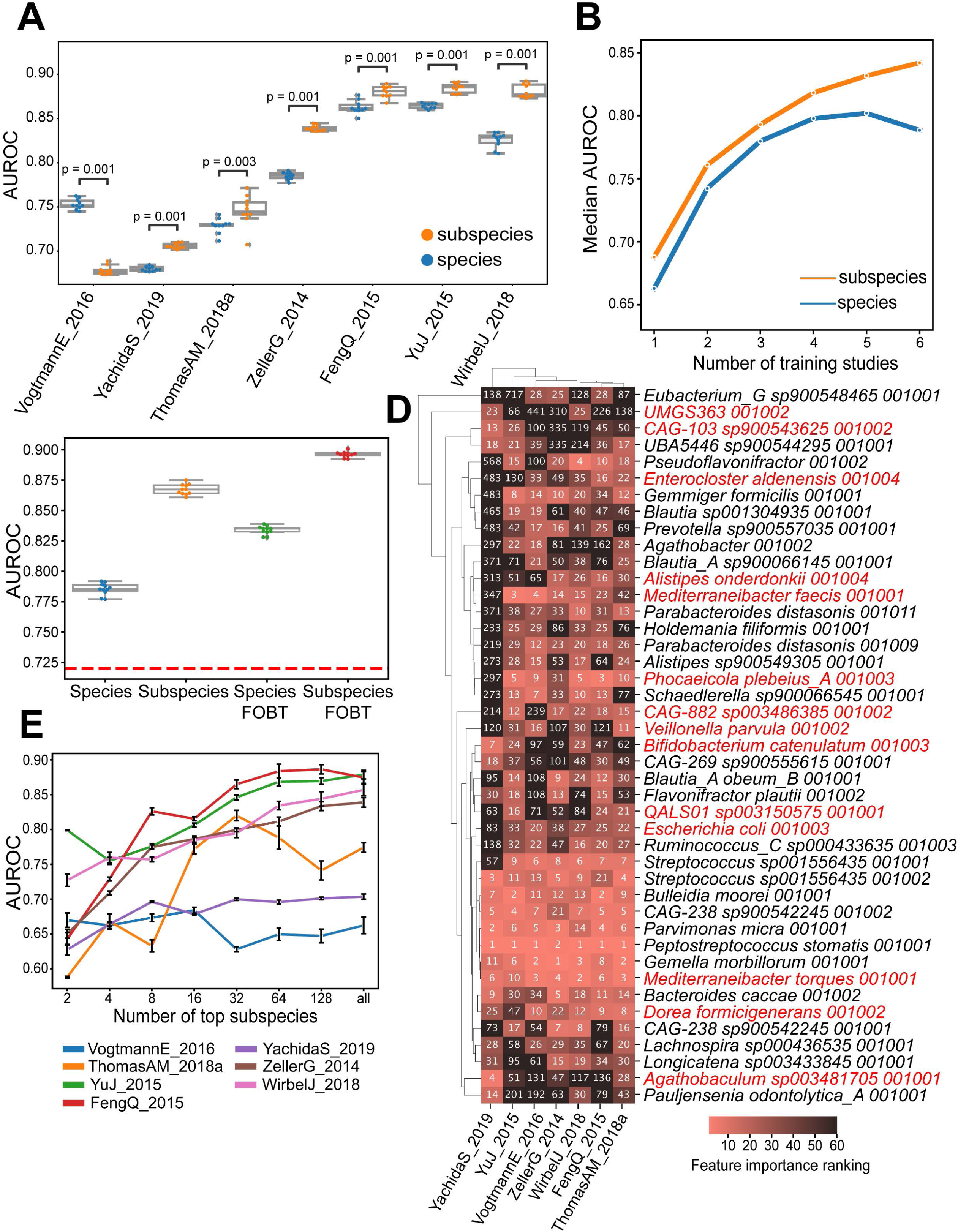
Subspecies are better predictors of colorectal cancer. (A) Boxplots of the AUROC at the species and subspecies level obtained through the LODO approach initialized ten times with different random states. The study used for testing is denoted on the x-axis, while all others were used for training. Difference in performance between species and subspecies level models was tested using Mann-Whitney U-test. (B) Line plot of median AUROC performances between different numbers of studies used for training at species and subspecies level. Each point represents a median AUROC for all combinations of that number of studies used for training, averaged for all test studies. Each model was initialized ten times. (C) Boxplots of the AUROC at species and subspecies level, with and without the result of FOBT used to weigh the model output. The red dashed line indicates the AUROC performance of FOBT alone. (D) Clustered heatmap showing the importance ranking of subspecies found as the top 10 most important ones for at least one study. Marked in red are subspecies whose sibling subspecies exist in the dataset but weren’t marked in the top 50. (E) Line plot of AUROC performance obtained through LODO approach, where only the most important subspecies are included during training. To prevent overfitting, subspecies were ranked using models not trained on the dataset used for testing.

Next, we investigated if the performance of the classifiers would increase with more data being available for training. For that, we trained the model on a different number of datasets using the LODO approach and found that the subspecies level didn’t reach the plateau, while the species level started decreasing in performance with datasets used (Figure 4B). The decrease in prediction performance could be a consequence of lower reproducibility at the species level demonstrated using univariate statistics (Figure S4D). Additionally, the difference between subspecies- and species-level performance persisted along the different training sizes, and a similar trend was observed when investigating per-study performance (Figure S5D). To address the possibility of further increasing the predictive power by incorporating additional parameters, we included results of the fecal occult blood test (FOBT) available for study ZellerG_2014^41^. Since this information isn’t available in other datasets, we used it to weigh the final model output (see Methods for details). There, we achieved higher performance (median AUROC=0.893) compared to all other input combinations (Figure 4C), suggesting that inclusion of additional parameters can further improve the predictive performance.

To gain further insights into the improved performance of classifiers relying on subspecies-compared to the species-level ones, we ranked the importance of the individual subspecies learned by the trained models during the LODO approach, and compared them between the studies (Figure 4D). 14 out of 47 total top-ranked subspecies (29.78%, marked in red) didn’t have their sibling subspecies ranked in the top 50 of any of the studies. This suggests that the differential abundance of sibling subspecies, not quantifiable at the species level, enables classifiers to extract a CRC-related microbial signature with improved specificity. Therefore, we trained new models using only the top-ranked subspecies to define the minimal microbial signature required for successful training. We found that, for almost all studies, there was a plateau in performance at 64 top subspecies (Figure 4E), which can render classifiers more suitable and easier to interpret for diagnostic use in a clinical setting.

### Sibling subspecies can differently affect host physiology

The existence of closely related groups of bacterial genomes, some of which are associated with a certain host phenotype while others are not, presents a unique opportunity to investigate potential functional mechanisms of the bacteria in host physiology. To investigate whether indeed the subspecies resolution could provide insights into the underlying mechanistic origins of the subspecies-phenotype associations, we looked at the annotated gene differences between the subspecies. We considered both potential gain and loss of function: either by presence or absence of a complete gene, or by a destabilizing high-impact variant in one of the subspecies, effectively rendering that gene incapable of performing a specific function. Since subspecies groups in the HuMSub catalogue were initially defined based on variations in their gene sequences, we extracted this information by profiling the top ten subspecies positively associated with CRC (Figure 3C) together with their non-associated sibling subspecies and defining their functional differences. In accordance with preserving their evolutionary fitness, most of the detected SNVs didn’t have a significant predicted effect on protein stability (Figure S6A). However, there were many more cases of differences by high-impact (abs(ΔΔG) > 1) gene variants, than by the subspecies-specific accessory genes (Figure 5A). These high-impact variants have been attributed to genes involved in biosynthetic pathways of nucleotides and amino acids (Figure 5B). Interestingly, we uncovered differences in the type of the modules inactivated by the two mechanisms: nucleotide and amino acid biosynthesis were mainly affected by high-impact SNVs (Figure S6B), while gene specificity was responsible for differences in the synthesis of other types of molecules, such as metabolites and vitamins (Figure S6C).

**Figure 5:**
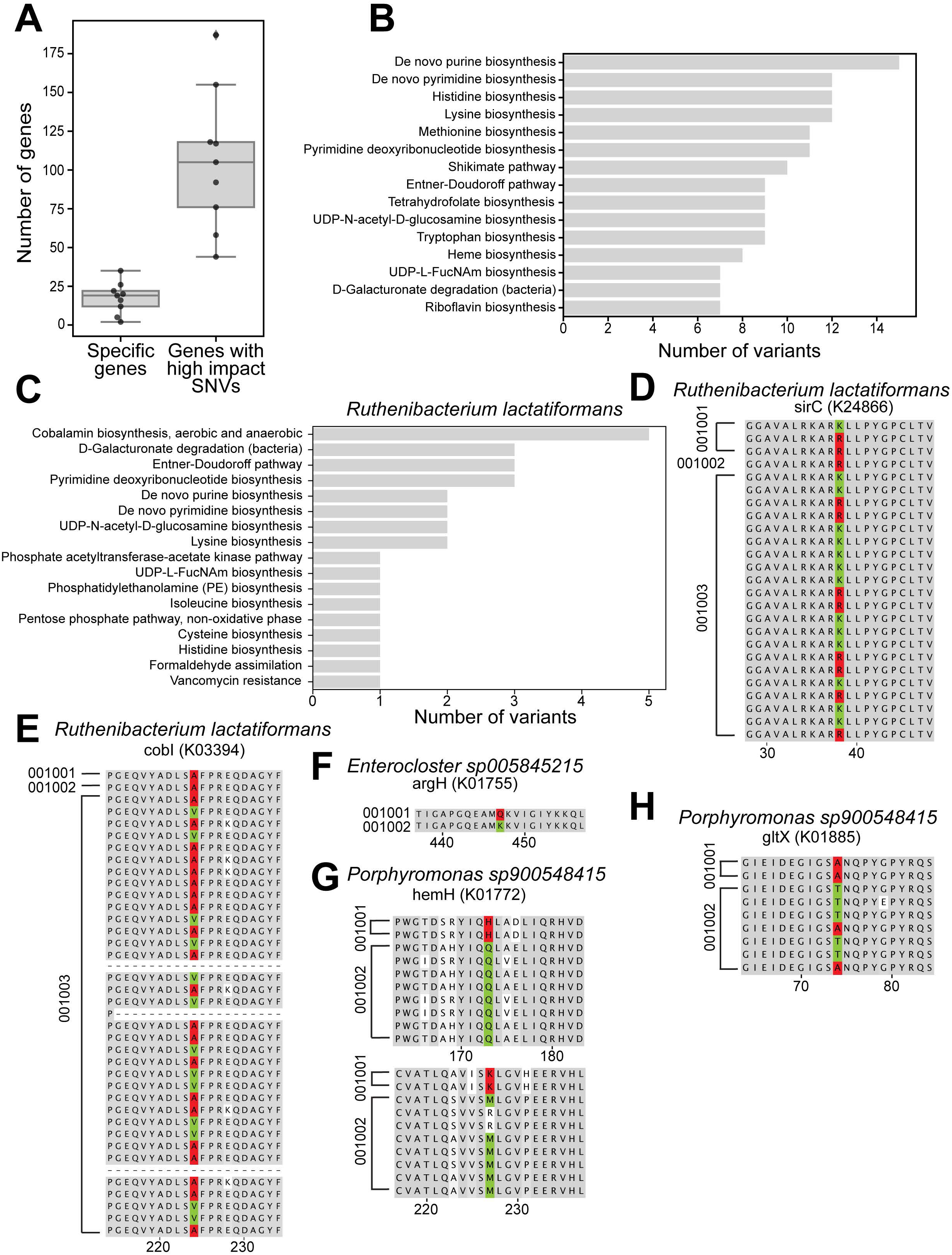
Sibling subspecies can differently affect host physiology. (A) Boxplot showing number of genes fully specific for a certain subspecies and number of genes with high-impact variants specific for a subspecies of top ten species associated with CRC. (B) Barplot denoting KEGG modules to which of the genes shown in A) they belong. (C) Barplot showing KEGG modules to which genes with high-impact variants of *Ruthenibacterium lactatiformans* belong. Aerobic and anaerobic cobalamin biosynthesis are grouped together. (D-H) MSA output of genes *sirC* (D)*, cobI* (E), *argH* (F), *hemH* (G) and *gltX* (H) found in the corresponding species. Red color marks variants that are predicted to lower the stability of the protein (abs(ΔΔG)>1). Sequences were dereplicated at the full length at the subspecies level. MSA visualizations are shortened to the region of interest.

We then looked into the possible role of several specific subspecies in the development of CRC. We first looked at *Ruthenibacterium lactatiformans 001003* as the top associated subspecies where its parental species is not associated with CRC (Figure 3C and 3D). Between the CRC-associated 001003 and its sibling subspecies *001001* and *001002*, we detected 183 high-impact variants of which 31 were located in 24 genes with annotated KEGG modules (Table S3). Five of those variants were found in four genes (K03394, K24866, K05936 and K02189) belonging to modules for aerobic (M00925) and anaerobic (M00924) cobalamin (vitamin B_12_) biosynthesis (Figure 5C). We found a highly destabilizing variant (abs(ΔΔG) = 2.4) located at position 38 of gene *sirC* (K24866) in all protein sequences of *001002* and some *001001* subspecies, which was absent in 80% of *001003* sequences (Figure 5D). Similarly, another high-impact destabilizing variant (abs(ΔΔG) = 2.7) was located at position 224 of gene *cobI* (K03394) in all sequences of non-associated *001001* and *001002* subspecies, not present in over 80% of *001003* sequences (Figure 5E). An additional disruptive variant (abs(ΔΔG) = 1.65) at position 5 of gene *cobM* (K05936) was found in all sequences of the subspecies *001001* (Figure S6D), and finally, two destabilizing variants were found at positions 195 (abs(ΔΔG) = 1.03) and 252 (abs(ΔΔG) = 1.0) of gene *cbiG* (K02189) of several *001001* sequences (Figure S6E). These data indicate that unlike the CRC-associated *001003*, the non-associated subspecies *001001* and *001002* have lost the ability to produce vitamin B_12_ in aerobic and anaerobic conditions due to these highly disruptive variants. Interestingly, higher concentrations of B_12_ are measured in the serum of CRC patients, and these B_12_ levels are positively associated with the stage of the cancer^53^. Higher B_12_ levels are associated with lower methylation levels of retrotransposable long interspersed nuclear elements (LINEs), both in tumor tissue and surrounding peripheral blood mononuclear cells^54^. LINEs are preventive biomarkers of CRC^55^. Supplementation of this vitamin increases the risk of CRC^56^, suggesting a potential causative effect of higher B_12_ levels on CRC progression.

We next investigated whether such mechanistic explanations are also observable with subspecies that show lower levels of association with CRC. We compared the functional potential of the *Enterocloster sp005845215* subspecies *001002* that was increased in CRC samples relative to its sibling non-associated subspecies *001001* (Figure S6F) and found 108 gene differences, from which 11 could be linked to 10 KEGG modules (Table S4). Among those genes, *argH* (KO K01755), which is a part of the biosynthetic pathway of the amino acid arginine (M00845), showed a highly destabilizing variant (abs(ΔΔG) = 1.3) at position 447 in the non-associated subspecies *001001* (Figure 5F). Interestingly, both asymmetric and symmetric dimethylarginines are higher in patients with CRC^57^. Moreover, increased dietary intake of arginine is associated with the increased incidence of CRC^58^, suggesting a causal relation between the increased arginine levels and CRC. Indeed, these previous findings led to development of arginine-degrading enzymes as a new class of anti-CRC drugs^59^. As another example, we compared subspecies *Porphyromonas sp900548415 001002* that was increased in CRC, to its CRC non-associated sibling subspecies *001001* (Figure S6G), and uncovered 130 different genes, 38 of which being annotated to 21 modules (Table S5). Among these, the gene *hemH* (KO K01772) that belongs to one of the pathways for the biosynthesis of heme (M00926), had two destabilizing variants in positions 173 (abs(ΔΔG) = 1.4) and 227 (abs(ΔΔG) = 2.0) within the non-associated subspecies *001001* (Figure 5G). Additionally, the gene *gltX* (KO K01885) was similarly destabilized (abs(ΔΔG) = 1.2) by a variant at position 74 (Figure 6H) in the same subspecies, which is part of another pathway of heme synthesis (M00121). Increased dietary intake, as well as the microbiota-produced heme, is linked to gut epithelial hyperproliferation^60^ and chronic intestinal inflammation that critically contribute to the development of CRC^61^, suggesting that the increased abundance of subspecies *001002* could be mechanistically linked, through heme production, to favorable conditions for CRC development. Together, these data indicate that at the subspecies resolution, microbiome profiling reveals an additional layer of information necessary for in-depth functional analysis, thus providing new insights into the underlying origins of microbiota-phenotype associations.

## Discussion

In this study, we report the HuMSub catalogue where we comprehensively define the human microbiota at subspecies resolution, and establish a panhashome-based method for rapid subspecies quantification and identification of specific genes that drive inter-subspecies phenotypic differences. Meta-analysis of colorectal cancer datasets reveals disease-associated subspecies that provide basis for development of machine-learning diagnostic algorithm that consistently outperformed species-level methods, emphasizing the significance of subspecies-level information in understanding microbiome-condition interactions.

Previous research, pioneered by efforts in a limited range of selected species focused on advancing from the classical species resolution, pointed to the need for a uniform and comprehensive reference system with subspecies that can be optimally compared across studies. To generate the HuMSub catalogue, we focused on predicting the coding sequences. In contrast to approaches using genome-wide SNVs, our method enables detecting consistent variations that cause actual phenotypic diversities. The sketching approach for subspecies delineation, as implemented in sourmash^26^, allowed us to define subspecies groups as collections of genomes with known sequences, which renders identifying the subspecies-specific functions relatively straightforward. With the custom subspecies-specific panhashome concept, we utilized the optimal computational efficiency of sketching to generate a highly compact resource that takes up only 127 MB of memory space, enabling its use without requiring extensive computational power, and facilitating easy sharing compared to the classical reference-based tools.

Even though higher taxonomic resolution could be followed by a loss of replicability across populations, the world-wide population analysis showed that HuMSub catalogue generalizes across highly diverse metagenomic samples, enabling its use in meta-analyses and other means of cross-cohort research. Moreover, HuMSub enables identification of functional microbiota differences relevant to the condition or the host phenotype, based on the differential sibling-subspecies associations and gene expression profile. Such approaches advance from the species-level methods currently used, since they allow accessing the functional changes and the direct contribution of the driver subspecies critical to a given phenotype in a direct comparison to their sibling, non-associated subspecies. On top of our investigation of subspecies associations on a global, population-level scale, it would be of interest in follow up studies to examine how they may influence each other within a single host. This could be further exploited by utilizing the workflows established here to extract subspecies-specific functions, following additional optimizations to increase its scalability. With that information, one could foresee generation of a computational metabolic model of the human gut microbiota more powerful than the state-of-the-art species-level ones.

Microbiota species-level machine learning approaches are promising as a new non-invasive tool for detection and diagnosis of certain diseases, including CRC. However, their direct applicability is still limited mainly by the insufficient prognostic rates embedded in the intrinsic level of information available at the species level. By using HuMSub, we found 204 CRC-associated subspecies, out of which, 95 (46.5%) were instances where at least one subspecies was significantly associated with CRC and at least one sibling subspecies was not, and we uncovered 26 subspecies associations, not detectable at the corresponding parental species level. Based on this new layer of identified associations between the microbiota and CRC, we found that irrespectively with or without any prior knowledge, subspecies-based classifiers outperformed the corresponding species-level models in distinguishing samples from healthy and diseased subjects.

Given its superior prognostic performance, training subspecies-based machine learning models by incorporating additional studies, and integrating them with simple pre-diagnostic tests, as demonstrated in this study, alongside modeling for various risk factors such as age, lifestyle, and medical history, could facilitate development of robust diagnostic approaches. These approaches, based on machine learning or deep learning techniques leveraging microbiota sequencing data, would enable accurate detection of diverse diseases, including CRC and other cancers, and prediction of treatment responders versus non-responders. HuMSub increases inter-study reproducibility and reduces variability between metagenomic studies, thereby enabling establishment of a consistent microbial signature for a host phenotype. Dwelling on the subspecies differences, HuMSub catalogue enables discovery of new mechanistic insights into the interplay between the microbiota and host phenotype, and sets the ground for analyses of new and reanalyses of existing datasets at an unprecedented depth.

### Limitations of the study

To query the biogeographical distribution using as many samples as possible, we utilized a pre-made sourmash index^32^. For that, we were restricted to a kmer size of 21, which limits false positives, but there is the chance that it may not detect all of the subspecies that were delineated in our study, possibly resulting in lower general prevalence (Figure 2A). Even though previous studies^17,33–37^ support our findings in context of the African populations, we can’t exclude that some subspecies could be even less geographically restricted. Moreover, since it is currently not feasible to perform the analysis with all generated hash values, the tools have to implement a filtering approach, such as scaling in sourmash. Even though empirically and experimentally tested, removal of hash values from a sketch inherently results in a loss of information. Although unlikely, we can’t exclude the possibility that the hash value scaling may translate to lower quantification power.

## Author contributions

MaT conceptualised and developed the study, wrote most of the code and wrote the first version of the manuscript. SK co-initiated the subspecies exploration and wrote most of the code regarding subspecies clustering. EZ co-supervised the work and advised on the methodologies. MT co-initiated, conceptualised, developed and supervised the work, and wrote the manuscript, with input from all co-authors. All co-authors read and approved the manuscript.

## Acknowledgments

We thank all members of our labs for discussions. The work was supported in part by the Clayton Foundation for Biomedical Research and the European Research Council (ERC) under the European Union’s Horizon 2020 research and innovation programme (ERC Consolidator Grant agreement No. 815962) to MT.

## Methods

### Resource availability

### Lead contact

Further information and requests for resources should be directed to and will be fulfilled by Mirko Trajkovski.

### Materials availability

This study did not generate new unique reagents. The study generated new computational tools made publicly available upon acceptance.

### Data and code availability

- This paper analyzes existing, publicly available data. These accession numbers for the datasets are listed in the key resources table.
- All original code is deposited at Zenodo and GitHub and it is made publicly available upon acceptance.
- Any additional information required to reanalyze the data reported in this paper is available from the lead contact upon request.

## Method details

### Genome filtering and the HumGut genome collection

The HumGut genomes are clustered by Hiseni et al.^21^ into species-level OTUs at 95% (HumGut_95) and 97.5% average nucleotide identity (ANI) clusters. Since some species could contain relatively diverse genomes, we used the 95% ANI clusters of the HumGut instead of direct GTDB and NCBI species for all further analyses.

We quality-filtered the HumGut collection using a custom workflow available at https://github.com/trajkovski-lab/Quality-filtering. Prodigal^62^ was run using the “-p meta” flag, which corresponds to the updated “anon” flag, used for metagenomes, low-quality draft genomes, small viruses, and small plasmids. The default output included all predicted genes, including the partial genes. The genes were then used as input for a two-fold filtering approach. The first fold was the detection of contaminated genomes, where we used GUNC^24^ with GTDB reference and kept only genomes marked as *‘passed’*. The second fold was the completeness and contamination assessment using BUSCO^25^. To best utilize the BUSCO approach, we performed a provisional run on a set of genomes containing one representative per HumGut_95 cluster in order to get their BUSCO taxonomic clade. Then, we used that information to run BUSCO on all genomes, now forcing the run on the previously selected taxonomies. This was done to prevent incomplete genomes from being evaluated at the lowest taxonomic clade, thus potentially obtaining an artificially high completeness score. The quality of analyzed genomes was assessed using the following formulas:

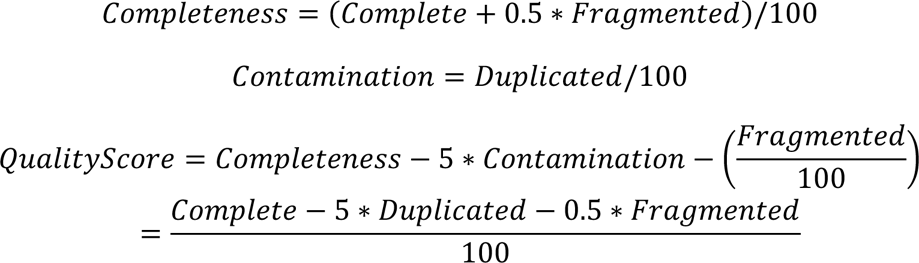

and only genomes with *QualityScore > 0.7* were kept for further use.

### Clustering

We relied on genes called during the quality-filtering step as the input for sketching using sourmash^26^ with the *“sketch”* command and parameters *“-p protein,k=51,scaled=1”*. The output of the *sketch* command for all genomes that passed QC and belonged to one HumGut_95 cluster was further processed together using the *sourmash compare* command with “--ksize 51 –protein--ani” flags. This command produces a similarity matrix, where values represent ANI estimates from protein sketches, which differs from the standard use of ANI. This similarity matrix was then transformed into a distance matrix (1 – similarity_matrix) and used for hierarchical clustering, where the input genomes were randomly attributed to training and testing sets. We then performed clustering on these sets using the single linkage method, generating linkage matrices that describe the cluster formations. For possible cluster numbers up to 15, we assigned cluster labels and calculated prediction strengths by comparing labels between training and testing sets. The optimal number of clusters was determined based on the mean Prediction Strength across 15 iterations, selecting the highest cluster number that exceeds a predefined cutoff of 0.8, as suggested in the original publication by Tibshirani and Walther^27^. The maximal number of clusters was data-driven, where out of 977 clustering events, there is a median of 2, a mean of 2.93 and a maximum of 12 subspecies. Finally, hierarchical clustering was applied to the entire distance matrix. We named subspecies as follows: *[cluster95]001[subspecies_number]*, where *cluster95* is the original species-level cluster ID from the HumGut catalogue and *subspecies_number* was attributed during clustering. If no subspecies were detected within the input species-level cluster, all genomes were labeled as *[cluster95]001001*.

### Definition of Subspecies-specific hashes

To generate the subspecies-specific panhashomes we relied on Sourmash^26^ combined with custom scripts. Specifically, we used *“sourmash sketch-p dna,k=51,scaled=1000”* on genomes annotated with subspecies information to obtain their sketches. Then, using a custom Rust script, we parsed all sketches per species-level cluster and found all hash values specific to each subspecies. To be selected, the hash values needed to be found in more than 20% of the respective subspecies’ genomes while not present in more than 5% of all others. Finally, the specific hash values were compiled into one signature per subspecies and then indexed with *sourmash index*.

We generated a custom workflow to leverage information about specific hashes. There, sequencing reads from input metagenomic samples are sketched using the same parameters used to generate subspecies-specific panhashomes. Then, we relied on the *gather*^63^ algorithm from sourmash to quantify subspecies-specific hash values from the index. The default sourmash output was then summarized to produce the final table of relative abundances.

### Quantification benchmark

We used CAMISIM^64^ to simulate ten metagenomic samples through the *Art* simulator^65^ with default error profiles. Each sample was simulated using the mean fragment size of 350 and SD=30, for a total of 3 GBp per sample. For samples simulated using HumGut genomes as a source, we randomly selected 15 genomes per subspecies or all genomes from a subspecies if it is represented with less than that. Then, we used CAMISIM to further randomly sample 1500 genomes from our selection to represent subspecies with a maximum of three genomes. We used log distribution for the abundance profiles, with a mean value of 1 and SD=2.75.

To further validate our quantification approach for real-world use, we simulated a new set of metagenomic samples, now using genomes uploaded to RefSeq since 01.01.2022, after the publication of HumGut. We obtained the list of all genomes from https://ftp.ncbi.nlm.nih.gov/genomes/refseq/assembly_summary_refseq.txt, as of 16.02.2024. Further, we filtered the list to keep only genomes with NCBI taxonomic IDs found in HumGut. Then, we ran our quality filtering workflow and kept genomes that passed GUNC combined with BUSCO QualityScore > 0.8, which kept 2432 genomes. By increasing the quality threshold, we maximized the number of included genomes but still ensured that the differences between the estimated and ground-truth abundances are caused by the quantification approach itself and not by the incompleteness and contamination of input MAGs. We annotated those genomes with subspecies information. There, we used the *sourmash search* command with the “--containment” flag to estimate the containment score between an input genome and the HuMSub catalogue. This created a containment estimate between each genome and all of the subspecies in the catalogue. Finally, we assigned genomes to those subspecies where they showed containment over 0.8 (80%). With this, we ended up with 324 genomes used for metagenome simulation. Finally, we simulated sequencing samples using the same parameters as for HumGut genomes, now including all input genomes.

In order to further give context to evaluation metrics, we ran MetaPhlAn4^1^ with the vOct22_CHOCOPhlAnSGB_202212.1 database on the same simulated samples. To match the abundance estimations with the ground truth, we looked at all abundances estimated at the species level and extracted their NCBI taxonomic IDs. We used that to match them to species-level taxonomic IDs from the input genomes while ignoring abundances estimated at levels higher than species. We summed abundances from our quantification to obtain species-level abundances across subspecies from the same species.

For evaluation metrics, we used F1-score and Euclidian (L2) distance. For all quantifications, we used an abundance threshold at 0.001 (0.1%), thus ignoring all taxa simulated at a level lower than that. The F1-score is calculated as a harmonic mean of the precision (the ratio of true positive calls and total positive calls) and recall (the ratio of true negative calls and total negative calls). On the other hand, we used L2 distance as a measure of discordance between ground truth and estimated abundances using the *scipy.spatial.distance.euclidean* function in Python. The tracking of used computational resources was done through the *benchmark* directive within the SnakeMake workflow. There, each sample was analyzed with MetaPhlAn4 within a single rule, while sourmash quantification consisted of two – one for sample sketching and the other for the *‘gather’* command. The total resources for sourmash quantification were summed between those rules. The computational time was found in the ‘s’ field, while the total memory required was in ‘max_uss’.

### Subspecies biogeographical distribution

We used Mastiff^32^, a sourmash-based tool, to analyze the geographic distribution of subspecies. It enables querying the pre-generated sourmash index consisting of public human metagenomic samples. As a result, a table consisting of containment estimates for each input query is returned. We used 0.2 as the containment threshold for inferring subspecies presence/absence and subsequent prevalence calculations, meaning that a metagenomic sample must contain at least 20% of subspecies-specific hashes to be declared positive for that subspecies. Since the index was generated with the parameters *“k=21,scaled=1000”*, we regenerated the subspecies-specific panhashomes to match those requirements. Since we effectively lowered the resolution of sketching, we were able to find specific hashes for 3038 subspecies. We used curatedMetagenomicData^66^ to obtain metadata of metagenomic samples. Here, we filtered samples and kept only ones originating from fecal samples from healthy adult individuals. Further, to reduce the noise from countries with fewer samples, we decided to keep only those with more than 30 samples in total. With this, we ended up with 5272 metagenomic samples from Europe, Asia, North America, and Africa. For prevalence calculations at the country level, we called each country subspecies positive if at least 10% of its samples were positive.

### Public metagenomic datasets

We downloaded sequencing data from seven datasets^39–45^ from seven countries in total. We filtered the samples and kept only ones labeled as “CRC” or “control” in the “study_condition” column of metadata tables obtained from curatedMetagenomicData^66^.

### Sequencing data preprocessing

We preprocessed all downloaded data using the ‘QC’ module from the ATLAS v2^67^ with default parameters. In short, using tools from the BBmap suite v37.78, sequencing reads were quality-trimmed, and contaminations from the human genome were filtered out.

### Machine learning

We used the Light Gradient Boosting Machine (LGBM) classifier^51^ implemented in the lightgbm Python library. We constructed the models with 500 estimators, using ‘gbdt’ as the boosting type, ‘binary_logloss’ as the loss function, 0.01 learning rate, and 11 leaves. Here, we used the same subspecies-level quantifications as for meta-analysis. However, we obtained species-level abundances using MetaPhlAn4 and vOct22_CHOCOPhlAnSGB_202212.1 database with default parameters. Before training, we filtered out all features found at the abundance <0.001% in less than 85% of samples, to ensure that we only keep the highly prevalent and not very low abundant ones. This filtering approach enabled us to reduce the “Course of dimensionality” and eliminate the irreproducible features found only in a subset of samples. This means that for the LODO approach, we filtered features separately for each combination of studies included within training. With these criteria, of all detected subspecies on average 63.5% were filtered out.

We used two approaches to estimate the performance of the models: on all datasets together and the Leave-One-Dataset-out (LODO) approach. For the first one, we combined all datasets together and used tenfold cross-validation. For the latter, we used the LODO approach to simulate the real-world use case - six datasets were used for training, while the seventh one was used for testing, which we repeated five times. For all of the approaches, we reinitiated the model and, where applicable, the cross-validation ten times with different random states. The importance of features was assessed using the build-it ‘feature_importances_’ method of the LBGM classifier. To include information from FOBT, we trained the model using the LODO approach but increased the output value by 0.2 if the test sample was positive in FOBT.

### Identification of subspecies-specific genes

To guarantee that we find genes that finally resulted in the differential abundance between subspecies, we used subspecies-specific panhashomes from the HuMSub as a starting point. First, we found the minimal set of input genomes where those hash values could be found. Then, we combined those genomes and used *sourmash signature kmers-k 51* to translate them into kmers. Next, we used prodigal-derived gene calls obtained during the initial quality-filtering step to find the corresponding complete sequences from which the specific kmers were extracted. Then, we used MMseqs2^68^ to cluster the extracted genes based on their sequence similarity with the threshold at 0.9. Further, we annotated those representatives using DRAM^69^. If two or more genes were annotated with the same function, we merged those clusters and considered them together. There could be two mutually exclusive scenarios: a gene as a whole could be defined as present in only one of the subspecies, or it could be present in multiple subspecies but with subspecies-specific variants. If the first possibility was true, then we reported the whole gene as subspecies-specific. Alternatively, we performed the multiple sequence alignments (*MSA*) with muscle^70^ and defined positions of variants found in more than 80% of certain subspecies and not in more than 20% of all other subspecies. Then, we used PROSTATA^71^, a transformer-based tool, to predict the effect of a variant on the protein stability. All variants with abs(ΔΔG) > 1 were considered as high-impact variants. For MSA visualization, we used pyMSAviz Python library.

### Quantification and statistical analysis

#### Subspecies biogeographical distribution

To calculate if a subspecies is significantly differently prevalent between countries, we used a robust approach based on the Chi-Square Test of Independence (implemented as *scipy.stats.chi2_contingency* in SciPy^72^ Python library), followed by the Benjamini-Hochberg procedure for adjusting *P*-values, effectively controlling the false discovery rate. If the adjusted *P*-value for a species was lower than 0.05, we used the post-hoc Tukey’s Honestly Significant Difference (HSD) test to identify which pairs of countries were indeed different in prevalence. To summarize the information obtained through statistical testing, we ought to define a single metric, geographical enrichment score (GES) that would enable us to distinguish geographically restricted subspecies. We defined it as GES_subsp_=[(abs(median_difference) + log(median(adjusted *P*-value) + 0.0001)) / 2]. Finally, we scaled the GES across subspecies to [0,1].

### Meta-analysis

We used relative abundances obtained through our quantification workflow and arcsine-square root-transformed them. To adjust the transformed abundances for BMI and age, we created linear models that were fitted to the transformed abundances of all samples within each study individually, adjusted for disease presence, age, and BMI (abundance ∼ study_condition + age + BMI). Then, we used them to predict adjusted mean transformed abundances per study, which further served as input. For statistics, we relied on the metafor^73^ R package. We used the *escalc* function to estimate the effect sizes for each species and subspecies separately by calculating standardized mean difference (*Cohen’s d*). Then, we used those estimates with the rma function to calculate random effect sizes and the corresponding *P*-values. Finally, those *P*-values were corrected for multiple testing using the Benjamini-Hochberg procedure with the cutoff at 0.1.

### Machine learning

On each taxonomic level, we generated models using different random states to minimize the effect of the random model initialization on the performance estimation. To compare model performances, we used the Mann-Whitney U test with the Benjamini-Hochberg procedure to correct for multiple testing where applicable.

## Supplementary figure legends

**Figure S1:**
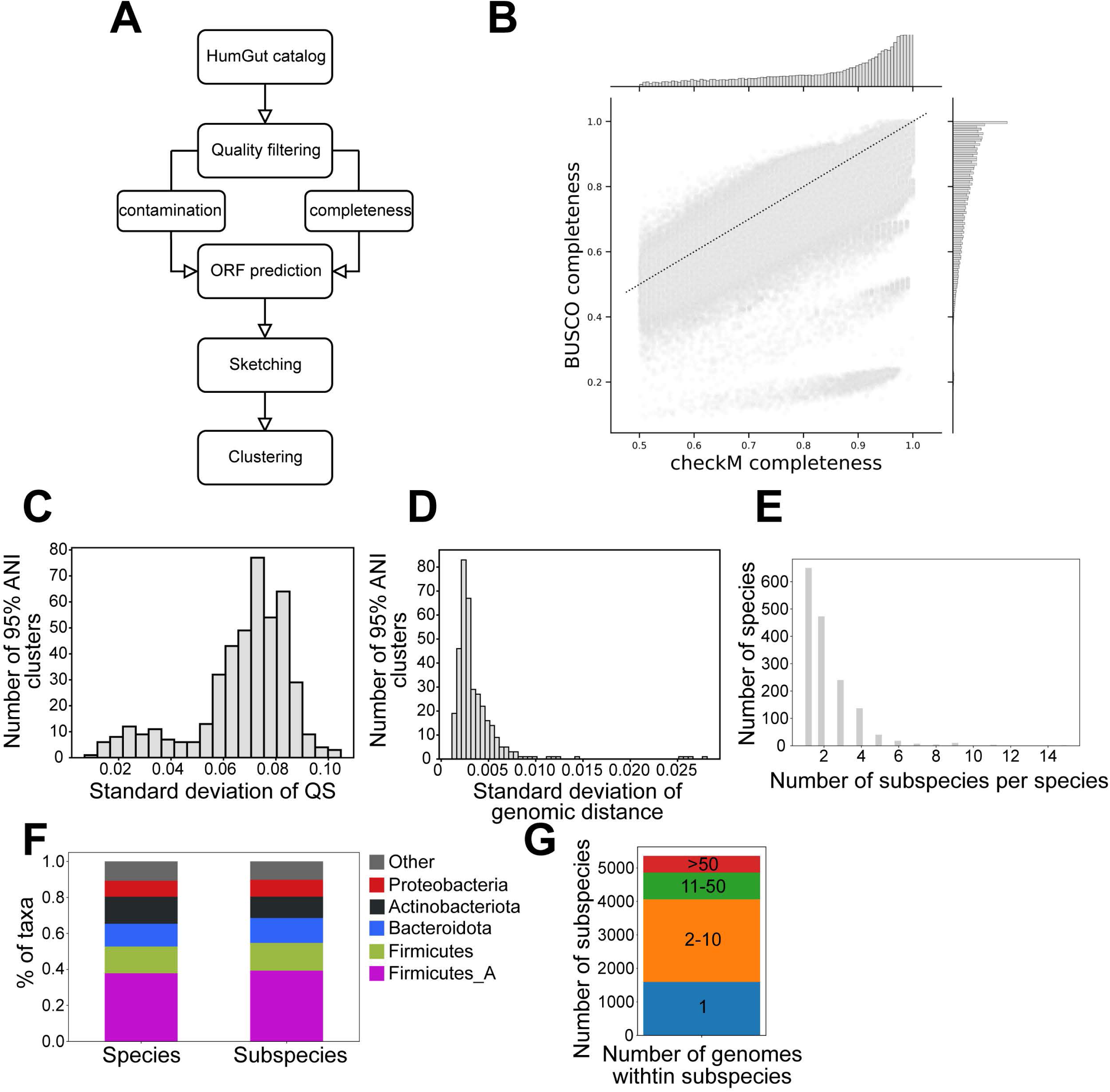
Summary of the subspecies delineation, related to Figure 1. (A) Workflow of subspecies delineation. Completeness estimation was done using BUSCO, while contamination estimation was done using both GUNC and BUSCO. (B) Jointplot of CheckM and BUSCO completeness assessments. CheckM completeness was obtained from the HumGut catalog. (C and D) Histogram illustrating distribution of standard deviation of BUSCO Quality score (C) and genomic distance (D) over 95% ANI clusters with more than 50 genomes. Genomic distance is the same one used for subspecies delineation. (E) Histogram showing the distribution of subspecies number per species. (F) Stacked barplot showing the percentage of species and subspecies grouped at the phylum level. (G) Stacked barplot describing the distribution of subspecies cluster sizes.

**Figure S2:**
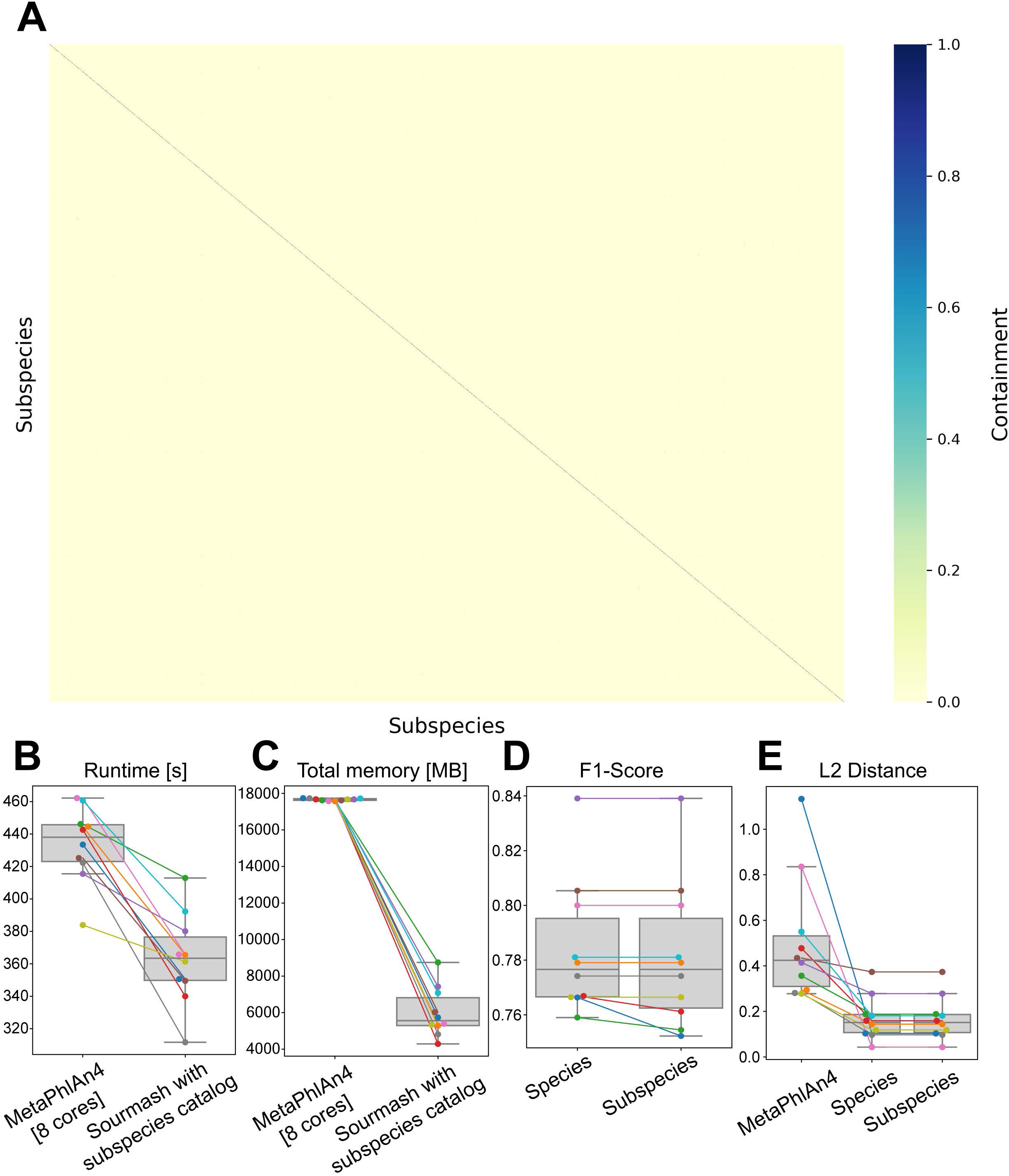
The subspecies-specific panhashome approach is efficient and precise in quantifying subspecies, related to Figure 1. (A) Heatmap showing pairwise-calculated containment of subspecies-specific hashes across all subspecies. Subspecies labels on the x- and y-axis are omitted for visualization purposes. (B) Boxplot denoting runtime for MetaPhlAn4 run on 8 cores using default options and sourmash-based subspecies quantification. (C) Boxplot denoting memory required for MetaPhlAn4 using default options on 8 cores and sourmash-based subspecies quantification. (D) Boxplot of F1-score distribution at species and subspecies level calculated on samples simulated using genomes outside of the HumGut catalog. (E) Boxplot of L2-distance distribution at species and subspecies level calculated on samples simulated using genomes outside of the HumGut catalog.

**Figure S3:**
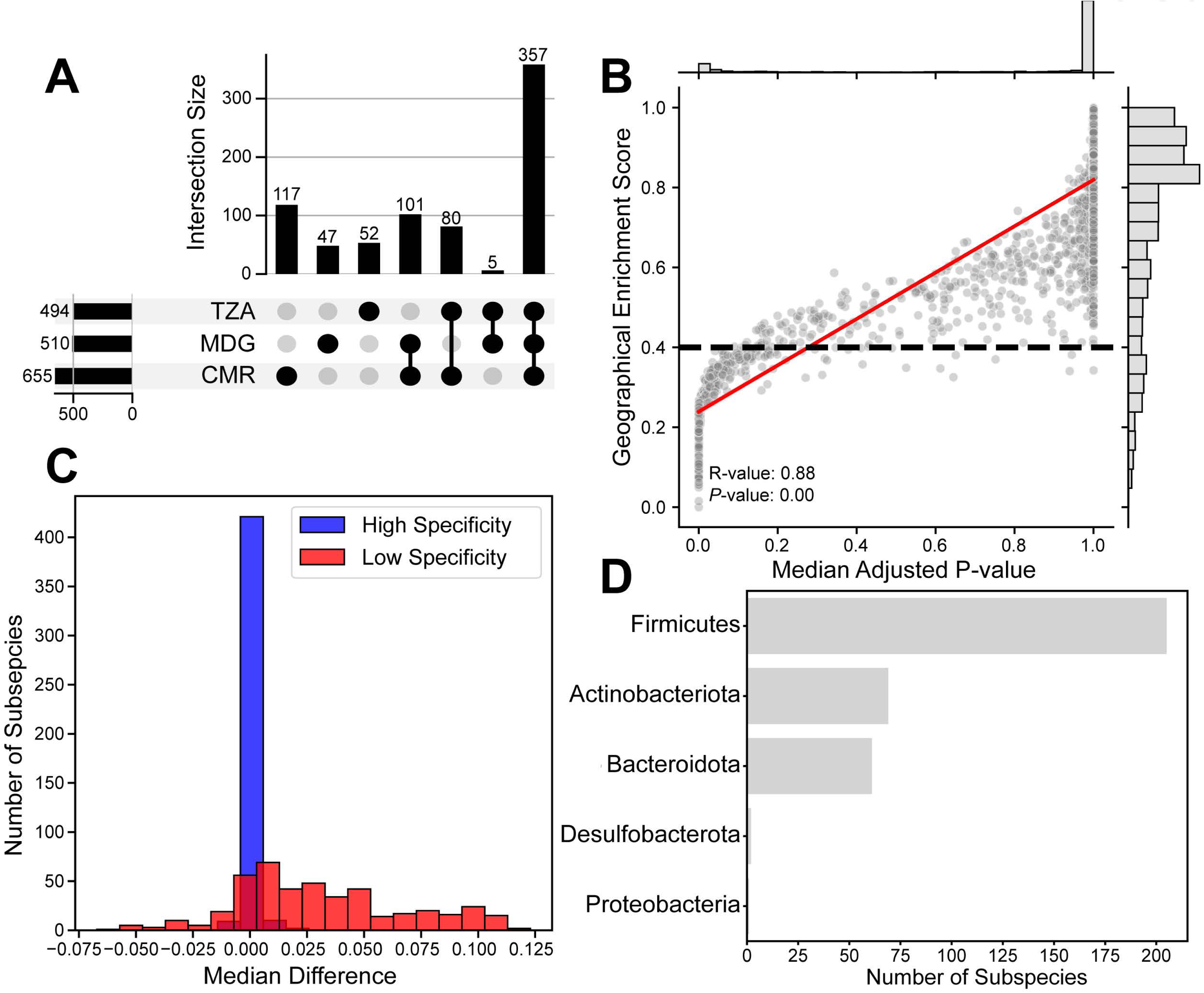
Geographical Enrichment Score (GES) correctly estimates subspecies’ geographical restriction, related to Figure 2. (A) UpSet plot showing the number of detected subspecies shared by different African countries. (B) Jointplot denoting the correlation between GES and median adjusted *P*-value obtained through pairwise country comparisons for each subspecies. (C) Histogram showing the distribution of median differences in prevalence between countries for each subspecies. In red, subspecies marked as “low specific” (GES < 0.6); in blue, subspecies marked as “high specific” (GES > 0.6). (D) Barplot showing a number of geographically restricted (GES > 0.6) subspecies grouped into phyla per NCBI taxonomy.

**Figure S4:**
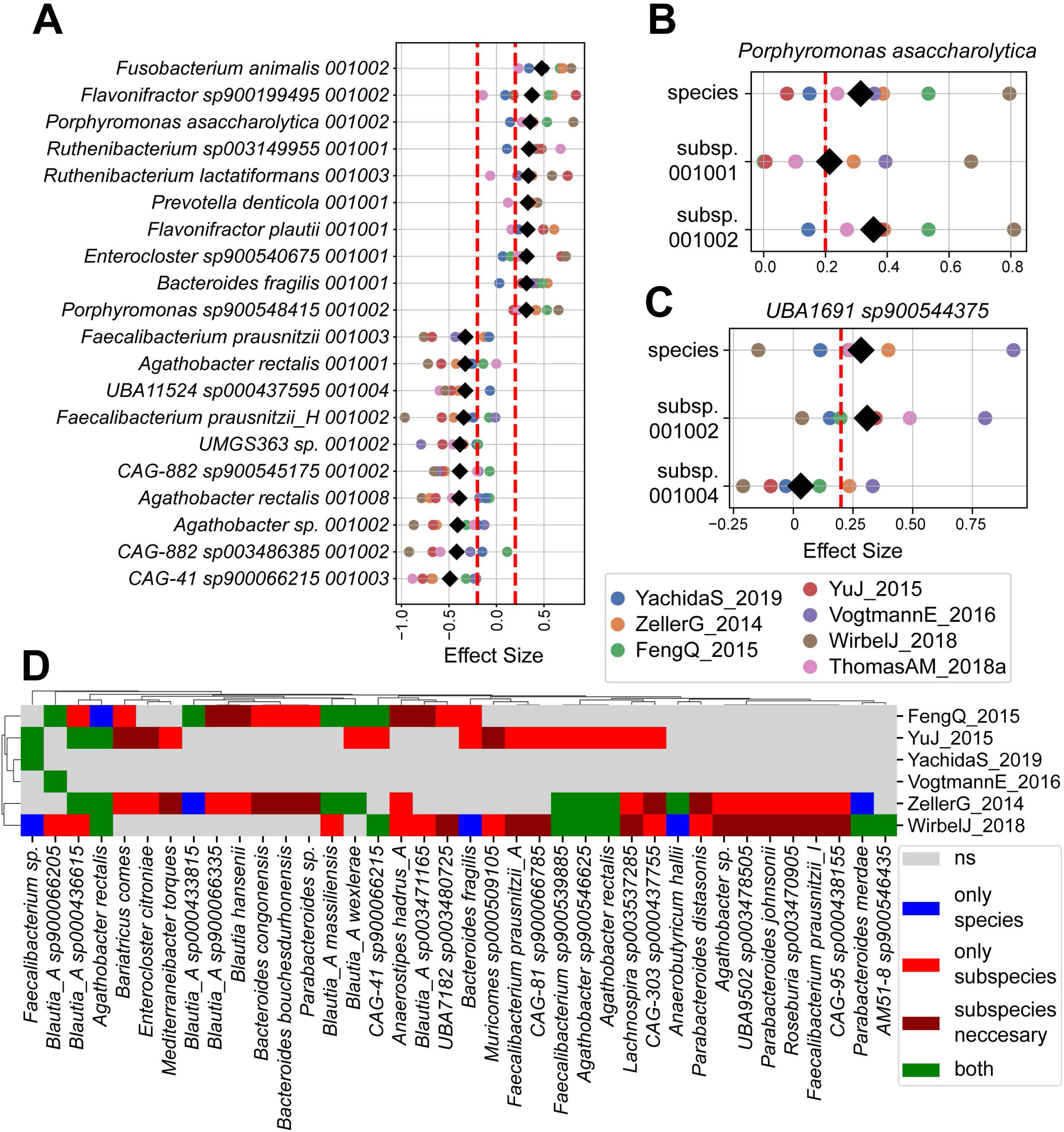
Sibling subspecies are differentially abundant between healthy and CRC samples, related to Figure 3. (A) Forest plot of effect sizes and the random effect sizes of the top ten subspecies positively and negatively associated with CRC, while at least one of their sibling subspecies isn’t (Blue dots from Figure 3A). (B and C). Forest plots of effect sizes and random effect sizes for *Porphyromonas asaccharolytica* (B) and *UBA1691 sp900544375* (C) species and their subspecies. Color dots represent effect sizes estimated for each study individually using standardized mean difference, while the black diamonds represent random effect sizes. (D) Clustered heatmap showing significantly associated species using univariate statistics on each study separately. At the subspecies level, if any of the subspecies are associated, the species is marked as associated. “Subspecies necessary” denotes associations where the subspecies level is necessary for a corresponding species to be associated in two studies. Species and subspecies are significantly associated if abs(log_2_FC) > 0.5 and FDR < 0.2.

**Figure S5:**
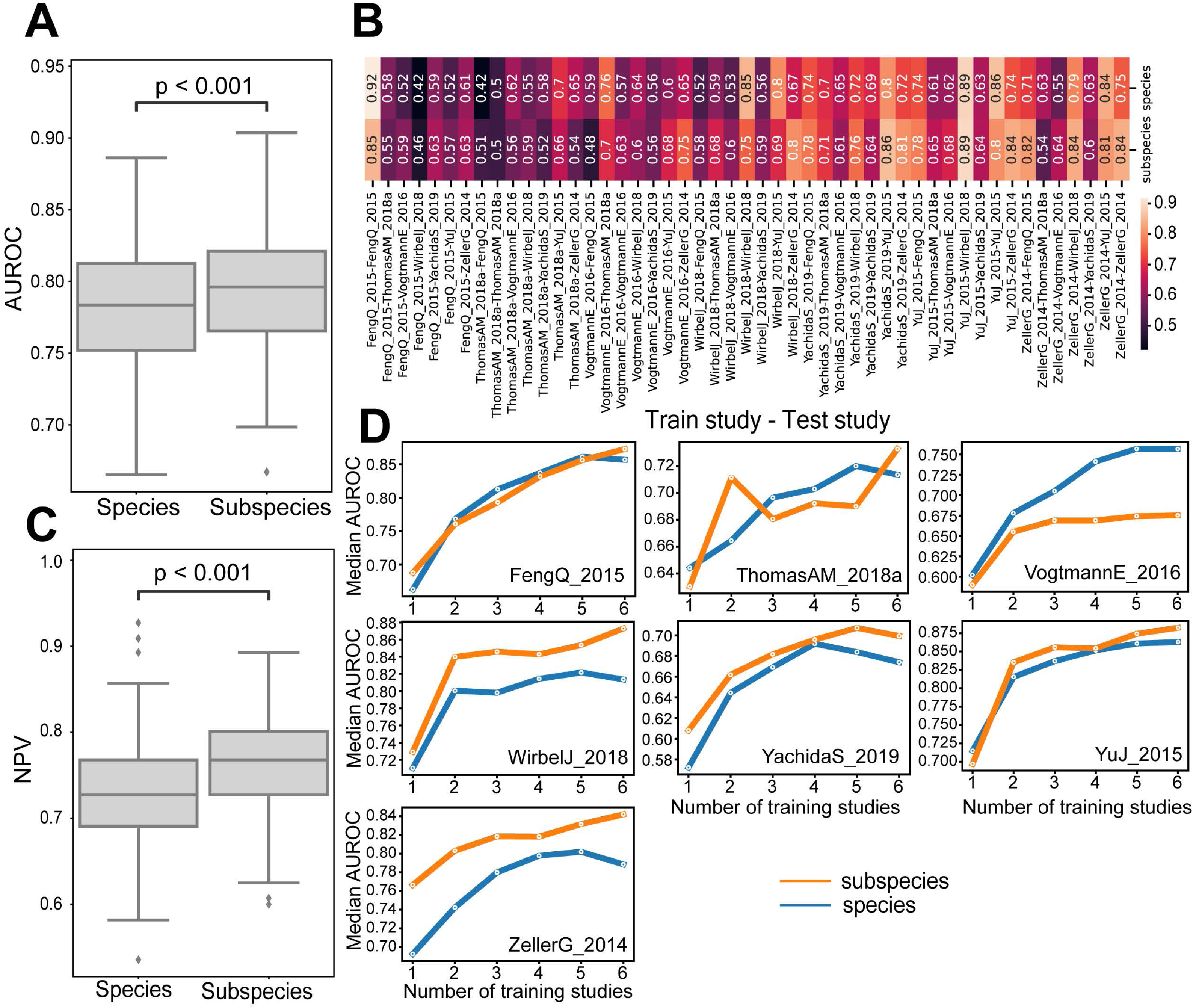
Subspecies enable improved prognostics of CRC, related to Figure 4. (A) Boxplots of the AUROC at the species and subspecies level obtained through the ten-fold cross-validation initialized thirty times with different random states. (B) Heatmap showing median AUROC of species- and subspecies-level models when trained and tested only on one dataset. (C) Boxplot showing the negative predictive value (NPV) of species- and subspecies-level models obtained through ten-fold cross-validation initialized thirty times with different random states. (D) Line plots of median AUROC performances between different numbers of studies used for training at species and subspecies level. Each point represents a median AUROC for all combinations of that number of studies used for training, where each model was initialized ten times. Indicated study refers to a study used for testing.

**Figure S6:**
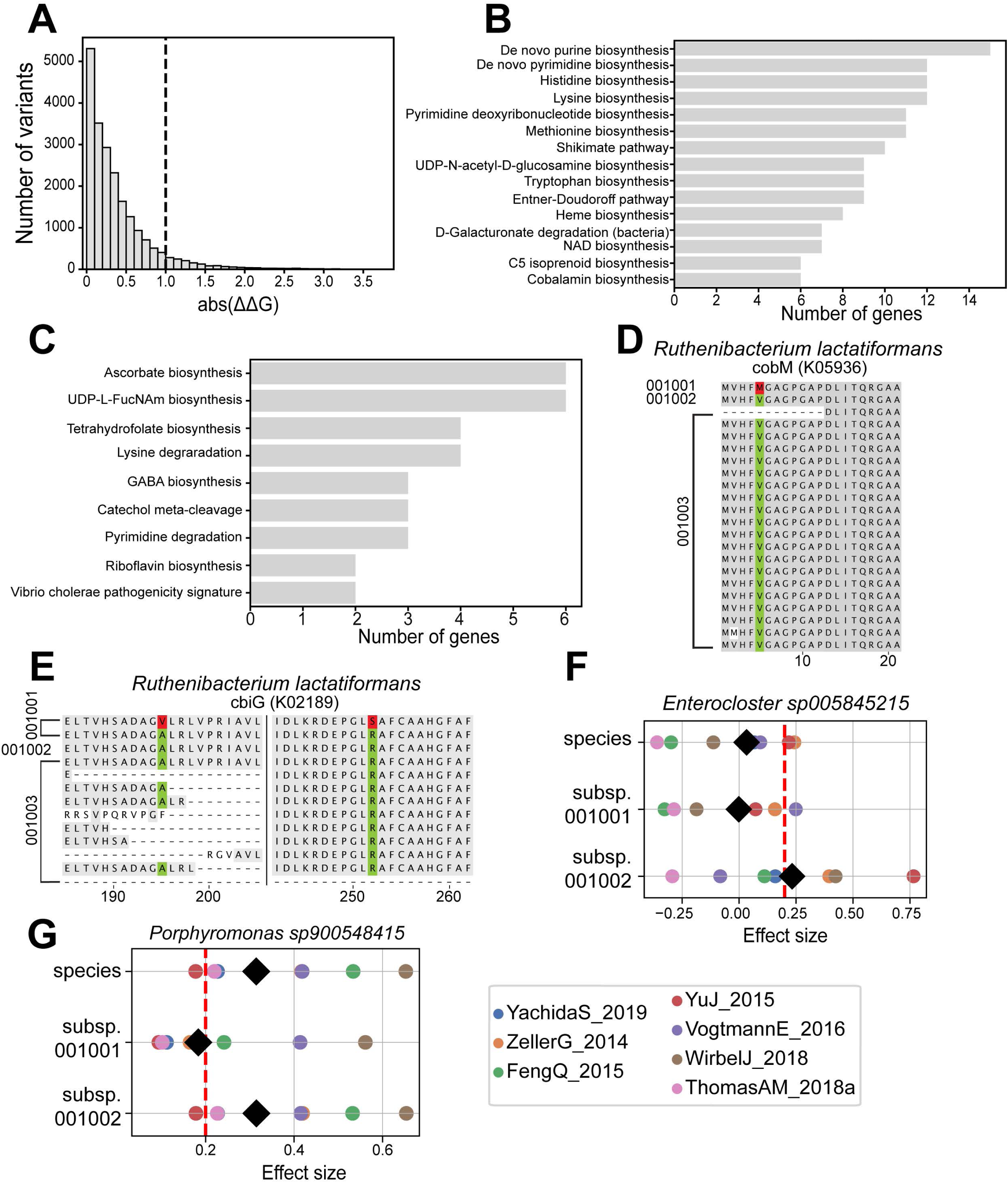
Subspecies-specific genes and variants, related to Figure 5. (A) Histogram showing the frequency of all effects of variants defined as subspecies-specific in the top ten subspecies associated with CRC. The vertical dashed line at abs(ΔΔG)=1 denotes the threshold for high-impact variants. (B) Barplot denoting KEGG modules to which genes that contain high-impact subspecies-specific variants belong. (C) Barplot denoting KEGG modules to which genes that are detected as fully subspecies-specific belong. (D and E) MSA output of genes *cobM* (D) and *cbiG* (E) from *Ruthenibacterium lactatiformans*. Red color marks variants that are predicted to lower the stability of the protein (abs(ΔΔG)>1). All protein sequences were dereplicated at the subspecies level. MSA visualizations are shortened to the region of interest. (F and G) Forest plots of effect sizes and random effect sizes for *Enterocloster sp005845215* (F) and *Porphyromonas sp900548415* (G) species and their subspecies. Color dots represent effect sizes estimated for each study individually using standardized mean difference, while black diamonds represent random effect sizes.

